# ProS-GNN: Predicting effects of mutations on protein stability using graph neural networks

**DOI:** 10.1101/2021.10.25.465658

**Authors:** Shuyu Wang, Hongzhou Tang, Peng Shan, Lei Zuo

## Abstract

**Motivation:** Predicting protein stability change upon variation through computational approach is a valuable tool to unveil the mechanisms of mutation-induced drug failure and help to develop immunotherapy strategies. However, some machine learning based methods tend to be overfitting on the training data or show anti-symmetric biases between direct and reverse mutations. Moreover, this field requires the methods to fully exploit the limited experimental data.

**Results:** Here we pioneered a deep graph neural network based method for predicting protein stability change upon mutation. After mutant part data extraction, the model encoded the molecular structure-property relationships using message passing and incorporated raw atom coordinates to enable spatial insights into the molecular systems. We trained the model using the S2648 and S3412 datasets, and tested on the S^sym^ and Myoglobin datasets. Compared to existing methods, our proposed method showed competitive high performance in data generalization and bias suppression with ultra-low time consumption. Furthermore, method was applied to predict the Pyrazinamide’s Gibbs free energy change for a real case study.

**Availability:** https://github.com/shuyu-wang/ProS-GNN.

## 1 Introduction

Proteins are composed of amino acids sequences, arranged into different groups. Changes to an amino acid due to DNA variation is called a missense mutation. Such mutations may result in changes of protein stability or protein misfolding[1]. One way to infer mutation induced protein stability change is to measure its ΔΔG. It refers to the change in folding energy between the mutant and wild state. The negative sign of ΔΔG indicates the variation decreases protein stability, and the positive sign means stability increases. Current study shows protein stability changes is one of the major underlying molecular mechanisms in multiple mutation-induced diseases[2]. Moreover, deeper insights into how specific mutations affect protein stability or interactions can identify possible drug resistance or sensitivity in patients[3], potentially leading to new precision medicines. This is especially important for genomic diseases such as cancers[4].

Recognizing the great potential of predicting protein stability changes upon mutation, researchers developed different computational tools, since they are low cost and high throughput. Prior attempts to predict protein stability were developed by molecular dynamics simulation[5]. They estimated free energy functions from protein structures using principles of statistical physics[6] or use knowledge-based terms of biophysical characteristics for regression fitting based on molecular mechanics[7, 8]. These methods showed advantages in characterizing the structural changes and the physical nature of the predicted folding free energy changes[9].

Data driven methods based on machine learning (ML)technologies are another promising branch to predict protein stability changes upon mutation. They are appealing due to efficient computation and high performance[10, 11]. Algorithms, such as support vector machines (SVM) [12–15], decision tree[16, 17], random forest (RF)[14, 18], gradient boosting[19, 20], and neural networks[21–23], or combinations of the above[24, 25] have been used for the purpose with preliminary successes. Before feeding the data to the ML pipeline, these methods need feature extraction. Some of the works only need sequence-based data and others might require structure-based data[26, 27]. Typically, methods using the 3D structures outperform the sequence-based methods[28]. Yet many prior works are prone to be biased on one direction variation or be overfitted[10]for practical usages. So the problem lies in how to fully exploit the limited experimental data and capture informative features.

End-to-end deep learning frameworks appeared to be a promising solution, which can enable useful features learning for various symbolic data. It can learn input features in the training process instead of fixing them and obtain data-driven features by directly utilizing the training dataset[29]. 3D CNN has been used to predict protein stability change with high performance[30]. It treated protein structures as if they were 3D images with voxels parameterized using atom biophysical properties. However, 3D grid representation entails void space where no atoms reside, leading to inefficient computation[31].

From the point of view of a many-body system, proteins are graphs by nature, in which a vertex is an atom and edges are a chemical bonds[32]. The total free energy is a sum over all atomic energy contributions. The graph neural networks(GNN) offers a new viable solution to elucidate the structure-property relationship directly from protein structural data. It shows high accuracy with a relatively low computational cost due to less parameters. It can identify important atom features determining molecular properties by analyzing relations between neighboring atoms[33], since the message passing process extracts structural features and then relates them with the target properties[34]. Similar approach has been demonstrated for atomic energies prediction from a quantum-chemical view[35], which implies GNN’s potential for protein related energy prediction. However, to the authors’ best knowledge, protein stability change prediction using GNN remained unexplored.

In this regard, we propose a novel and agile approach to predict protein stability change upon mutation using a deep learning model. After trimming the non-mutant part of the protein, the model maps the 3D structural information and element compositions of the protein to a high-dimensional representation, and automatically captures the key factors leveraging a gated GNN. Our key conceptual advance is implementing the model to predict structure-property following the underlying biochemistry law. We then demonstrate the method’s high performance by training with the S2648 and S3412 dataset, and tested on S^sym^ and Myoglobin dataset. Then we applied the method to predict the mutation effect on the drug for tuberculosis, Pyrazinamide (PZA), for drug resistance management.

## 2 Methods

### 2.1 Problem formulation

Our task is to predict the change of Gibbs free energy between mutant protein and wild protein. Our concept is derived from the many body Hamiltonian concept to embrace the principles of biochemistry, while maintaining the flexibility of a complex data-driven learning machine.

The input data is extracted from the PDB files, which contain the element composition and structural information. We formulate the protein feature vector ***h**_i_* for the ***i***th atom, containing the vertices and edges information. This feature vector encodes the element information, the number of adjacent atoms, the number of adjacent hydrogen atoms, implicit valence, and aromatic bonds. Thee feature vectors are combined to form the input feature matrix ***H*** = [***h*_1_**,…, ***h***_*n*+*r*_] for a protein, and 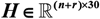. ***n*** and ***r*** are the atomic numbers of mutant and non-mutant parts, respectively. The wild and mutant type’s feature matrixes are denoted as 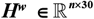 and 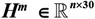. Similarly, the coordinate matrixes of the wild and mutant type atoms are 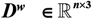 and 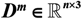, and the adjacency matrixes 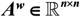 and 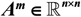 denote the adjacent relationship between atoms in the two proteins. These matrices form the input features to predict ***y***, ΔΔG. So the overall problem can be elucidated simply as:

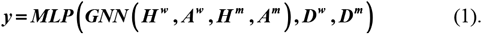

Here ***MLP*** is short for multiple layer processing.

### 2.2 Non-mutant part trimming

Different from prior methods, which process the complete protein feature matrix ***H***, our model only processes the information of the mutant residue and its two adjacent ones, denoted as 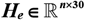. The residual 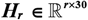 is trimmed by an automated script ( ***H*** =[<u>***H_e_ H_r_***]</u>).

### 2.3 Gated GNN

We expand the dimension of the feature vectors at the first layer 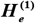 with a parameter matrix 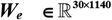 as 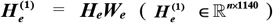. In the GNN, the *l*-th layer’s atom feature 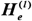 are processed by iterations of graph convolution to produce a set of updated atom features:

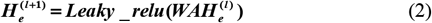

where 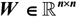 is a learnable weight matrix and 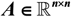. ***Leaky_relu*** is the activation function.

This is a simplified the message passing process, where each atom gathers local information from its neighboring atoms and bonds, and then update information. Through information sharing between atoms, a global feature can be extracted based on this technique. In this way, the GNN implicitly learns the property to be predicted from the structure.

To improve the feature extraction performance, we integrated the gating mechanism into the network[31]. The gated graph layer is a linear combination of 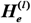 and 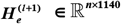:

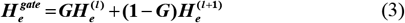

with

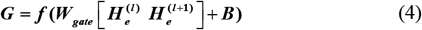

where 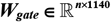 is a learnable matrix. 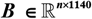 is a bias matrix. ***f*** is the Sigmoid non-linear activation function. The feature matrix of the gated connection 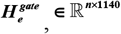 is then added to the first layer 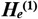:

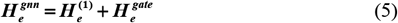

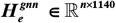 is the final output of GNN.

Then, we concatenate 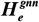 with the coordinate matrix 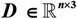 to generate a feature matrix 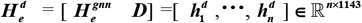. Here we assume 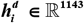 represents the energy contribution from the ***i***th atomic vector, so the sum of them corresponds to the total molecular energy:

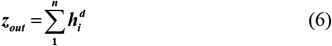

Finally, the feature vectors of the mutant type is subtracted from the wild type to get a contrasted feature vector. This feature vector is used for ΔΔG prediction after MLP. The process mentioned above can be expressed as follows:

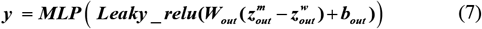

where 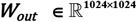 is a weight matrix and 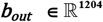 is a bias vector.

### 2.4 Training

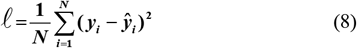

where ***y_i_*** and 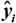 stand for the predicted ΔΔG and the experimental ΔΔG for the ***i***-th sample, respectively. Given all ΔΔG labels in the training data set when mutations occur, the training goal is to minimize the mean squared error (MSE) loss.

## 3 Experiments

### 3.1 Dataset

To train and test our model, we use several data sets listed below.

**S3421** contains 3421 experimentally determined mutations from 150 proteins.

**S2648** includes 2648 single-point mutation in 131 different globular proteins.

**S^sym^** contains 684 variations, and half of them are reverse variations.

**Myoglobin** is consisted of 134 mutations scattered throughout the protein chains. Myoglobin is a cytoplasmic globular protein that regulates cellular oxygen concentration.

### 3.2 Implement and evaluation

This work used a Nvidia Geforce GTX 3070 GPU for computing. We implemented the model using Pytorch and tuned the parameters by grid search. The dimension of the vertices vector is 1120 and the dimension of the full connected (FC) layers is 1024. The GNN and FC both have four layers. The model shows improved generalization, when the weight decay is 5×10^-5^, drop-out rate is 0.5 and the batch size is 16. In addition, the model is optimized by an Adam optimizer.

The following measures are adopted to evaluate the performance of ProS-GNN. The primary measures for evaluating prediction accuracy are the Pearson correlation coefficient (*r*) and the root-mean-squared error (RMSE) (*σ*) of the experimental and predicted ΔΔGs. ***r_dir-rev_*** and ***δ _dir-rev_*** are used to evaluate the prediction bias between the direct and reverse mutation prediction.

## 4 Results and Discussion

### 4.1 Trained using S2648 dataset

We first trained ProS-GNN using the S2648 dataset, and then tested with S2648, S^sym^, and Myoglobin datasets, respectively. When tested with the S2648 dataset, the proposed method achieved ***r*** = 0.62, ***σ*** = 1.11 on direct mutations (Fig. 2 (A)), ***r*** = 0.60, ***σ*** = 1.12 on reverse mutations (Fig. 2 (B)), and ***r*** = −0.94, ***δ*** = 0.04 on direct-reverse prediction(Fig. 2 (C)). The performance on the S^sym^ dataset achieved ***r*** = 0.61, ***σ*** =1.23 on the direct mutations, ***r*** = 0.56, ***σ*** = 1.30 on the reverse mutations and ***r*** = −0.94, ***δ*** = 0.04 on direct-reverse prediction (Fig. 2 (D)-(F)). Then, we compare our results with fifteen methods on the S^sym^ dataset and list them in Table1. It clearly showed our model outperformed other prior methods in prediction accuracy with little bias. On the Myoglobin dataset, the ProS-GNN achieved ***r*** = 0.48, ***σ*** = 1.27 on the direct mutations and achieved ***r*** = 0.43, ***σ*** = 1.19 on the reverse mutations and ***r =*** −0.90, ***δ*** = 0.07 on direct-reverse prediction (Fig.2(G)-(I)), which potentially suggested generalization in real-life applications. These results showed that our method could effectively learn feature representations with high performance.

**Fig. 1.**
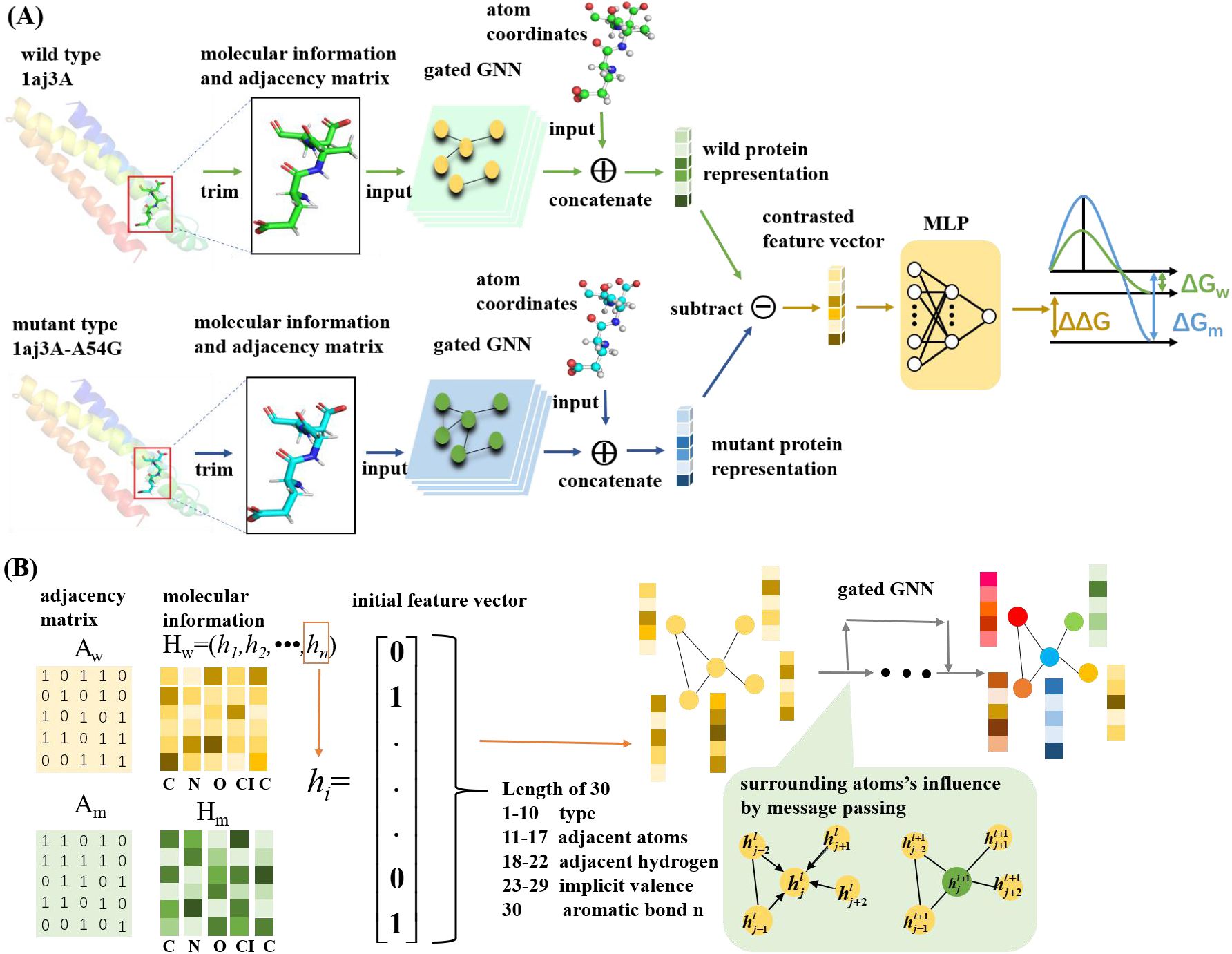
(A) The architecture of ProS-GNN. (B) Illustration of the input molecular features, adjacency matrix, and message passing in the gated GNN.

**Fig. 2.**
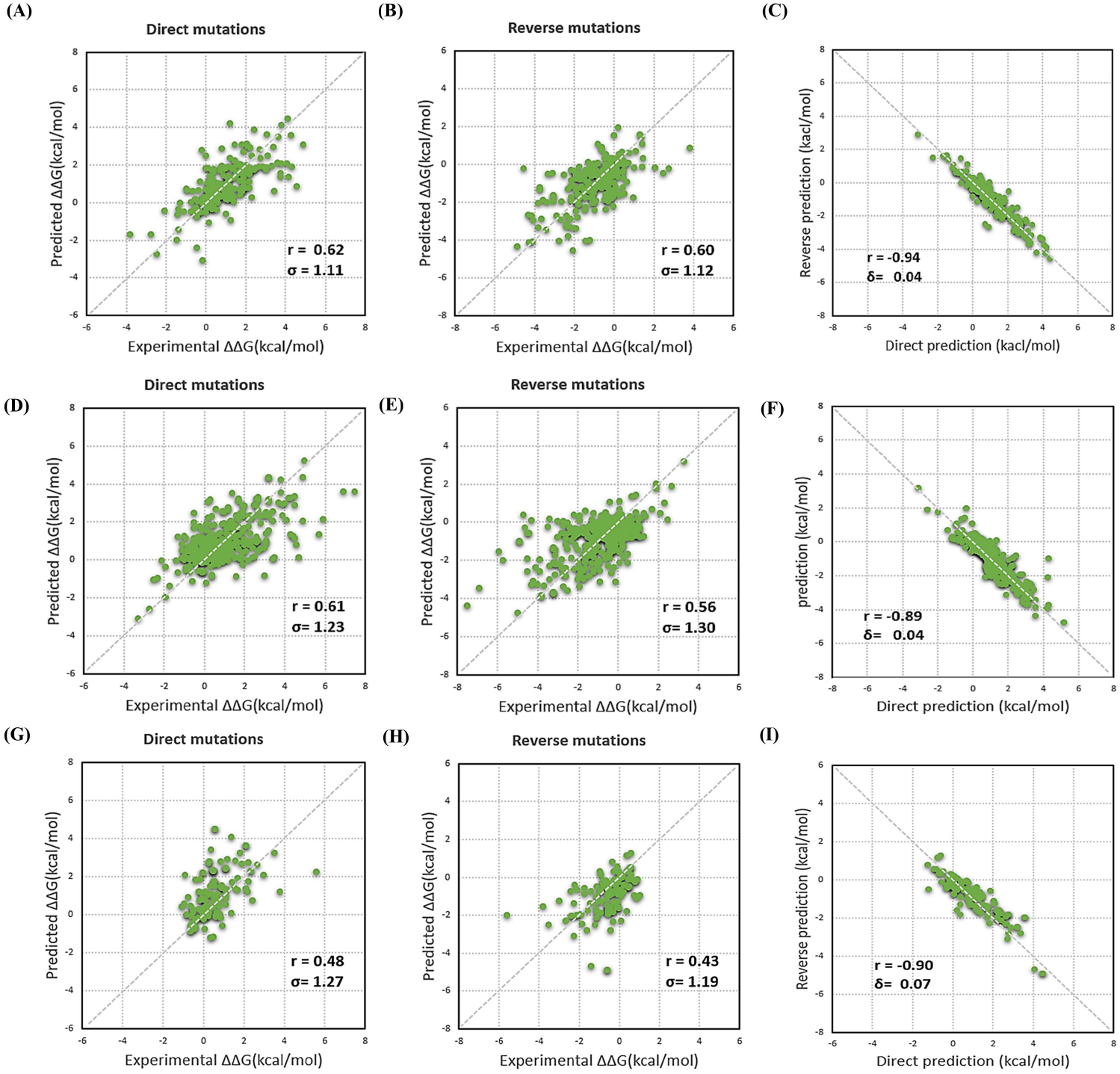
ProS-GNN trained using S2648 dataset and tested on the three datasets. (A) Predicting ΔΔG for direct mutations in S2648. (B) reverse mutations in S2648. (C) Direct versus reverse ΔΔG values of all the mutations in the S2648. (D) Predicting ΔΔG for direct mutations in S^sym^ (E) reverse mutations in S^sym^. (F) Direct versus reverse ΔΔG values of all the mutations in the S^sym^. (G)Predicting ΔΔG for direct mutations in Myoglobin. (H) reverse mutations in myoglobin. (I) Direct versus reverse ΔΔG values of all the mutations in the Myoglobin.

**Table 1.**
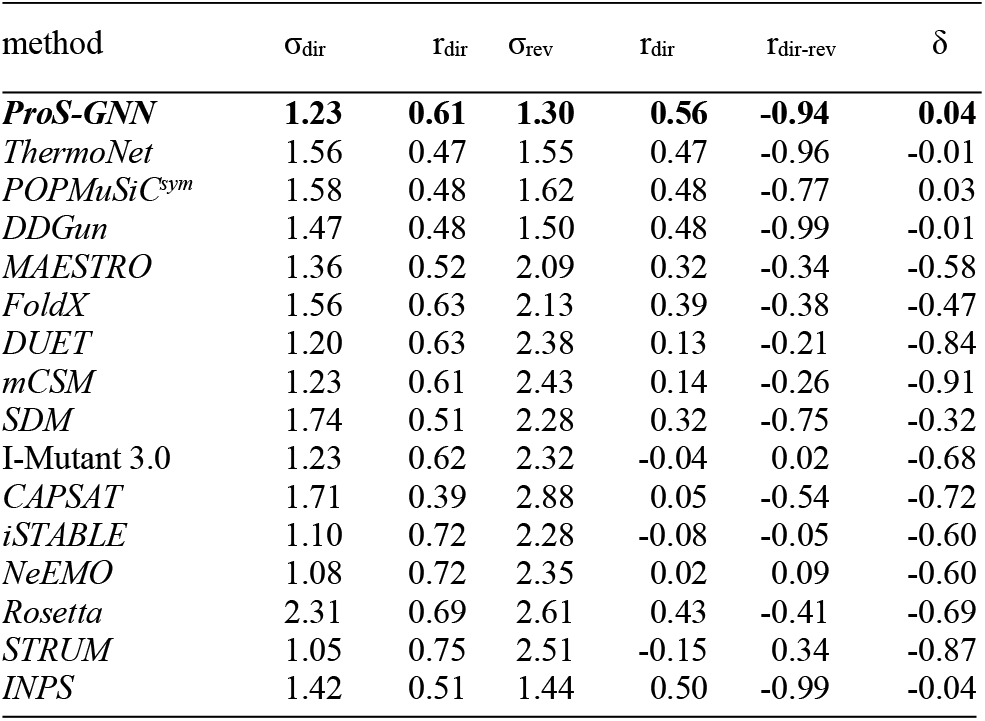
Comparison of different methods on the S^sym^ dataset.

### 4.2 Trained using S3421 dataset

Similarly, we also trained ProS-GNN using S3421 dataset and tested with S3421, S^sym^, and Myoglobin dataset, respectively. The testing results on the S3421 dataset showed ***r*** = 0.69, ***σ*** = 1.73 on direct mutations (Fig. 3 (A)), ***r*** = 0.71, ***σ*** =1.69 on reverse mutations (Fig. 3 (B)), and ***r*** = −0.95, ***δ*** = −0.21 on direct-reverse prediction(Fig. 3 (C)). When switched to the S^sym^ dataset, the ProS-GNN achieved ***r*** =0.51, ***σ*** = 1.47 on the direct mutations, ***r*** = 0.51, ***σ*** = 1.43 on the reverse mutations, and ***r*** = −0.95, ***δ*** = 0.21 on direct-reverse prediction (Fig. 3(D)-(F)). We found the performance was moderately inferior to the ones trained using S2648, which might be explained as S3214 shared no homology with S^sym^. Last, the tested performance on Myoglobin dataset was also competitive, as it achieved ***r*** =0.51, ***σ*** = 1.20 on the direct mutations prediction, ***r*** =0.45, ***σ*** = 1.13 on the reverse mutations prediction and ***r =*** −0.88, ***δ*** = −0.18 on direct-reverse prediction (Fig. 3 (G)-(I)).

**Fig. 3.**
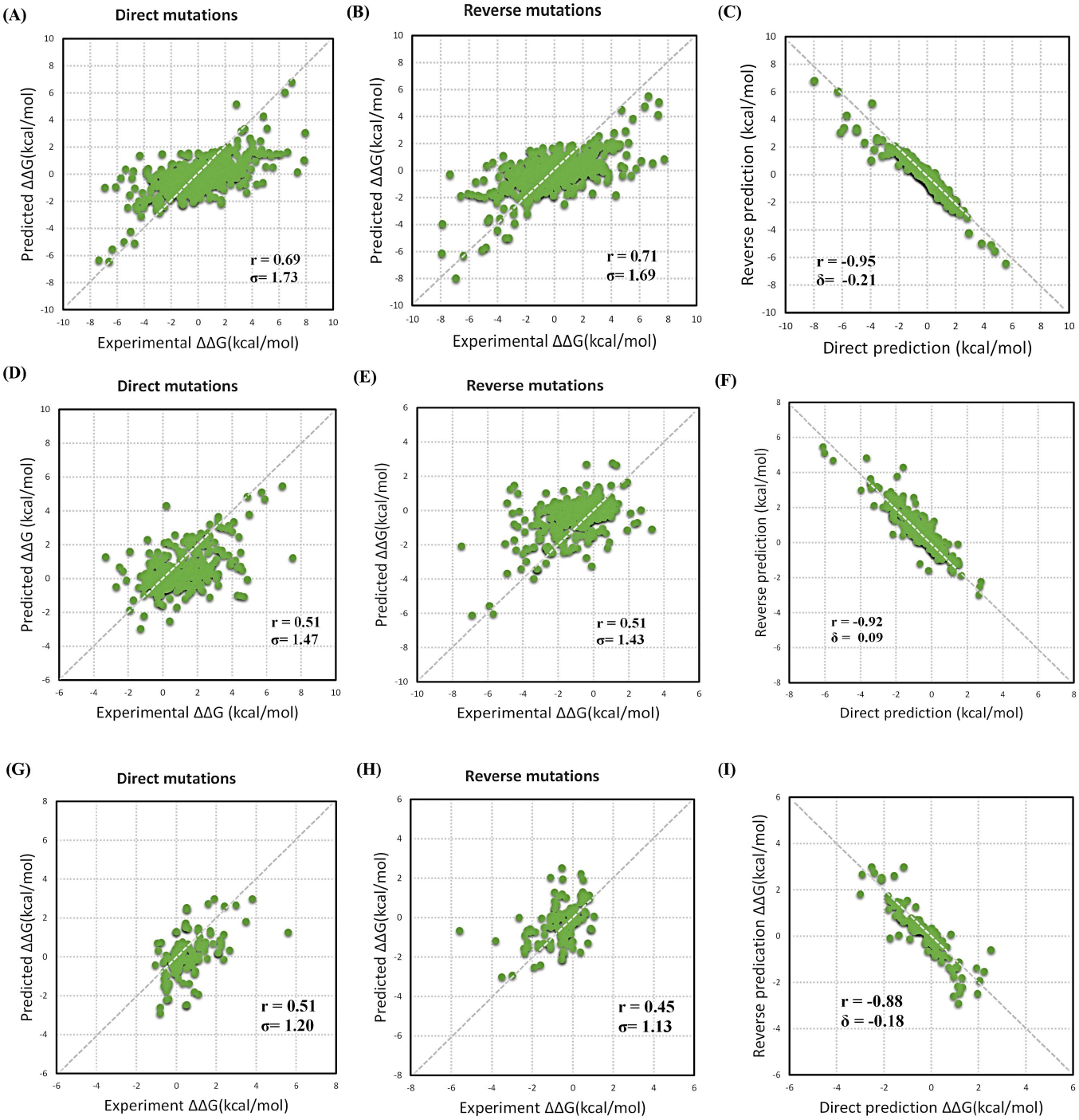
ProS-GNN trained using S3421 dataset and tested on the three datasets. (A) Predicting ΔΔG for direct mutations in S3421 (B) the reverse mutations in S3421. (C) Direct versus reverse ΔΔG values of all the mutations in the S3421. (D) Predicting ΔΔG for direct mutations in S^sym^ (E) reverse mutations in S^sym^. (F) Direct versus reverse ΔΔG values of all the mutations in the S^sym^. (G) Predicting ΔΔG for direct mutations in Myoglobin. (H) reverse mutations in myoglobin. (I) Direct versus reverse ΔΔG values of all the mutations in the Myoglobin.

### 4.3 Case studies: ribosomal protein S1(RpsA)

In the real case study, we used two single-point mutations, D343N and I351F, in Pyrazinamide (PZA). PZA is one of the first-line drugs, effective against latent Mycobacterium tuberculosis isolates. Resistance to this drug emerges due to mutations in pncA and rpsA genes, encoding pyrazinamidase (PZase) and ribosomal protein S1(RpsA), respectively.

We fetched the structure of RpsA (PDB ID 4NNI) from RCSB PDB, and generated the molecule structures of D343N and I351F with PYMOL(Fig. 4). The literature[36] indicates the ΔΔG of 4nniA-D343N mutation and 4nniA-I351F are 3.2 kcal/mol and 2.9 kcal/mol. Our proposed method predicted 1.6 kcal/mol and 2.6 kcal/mol, respectively. While the one of the results showed discrepancies between predicted and experimental value, the differences were within 1.6 kcal/mol, which was still normal considering the method’s RMSE. Therefore, it demonstrated the potential of ProS-GNN as a rapid estimator of ΔΔG upon protein mutations in a medical environment.

**Fig. 4.**
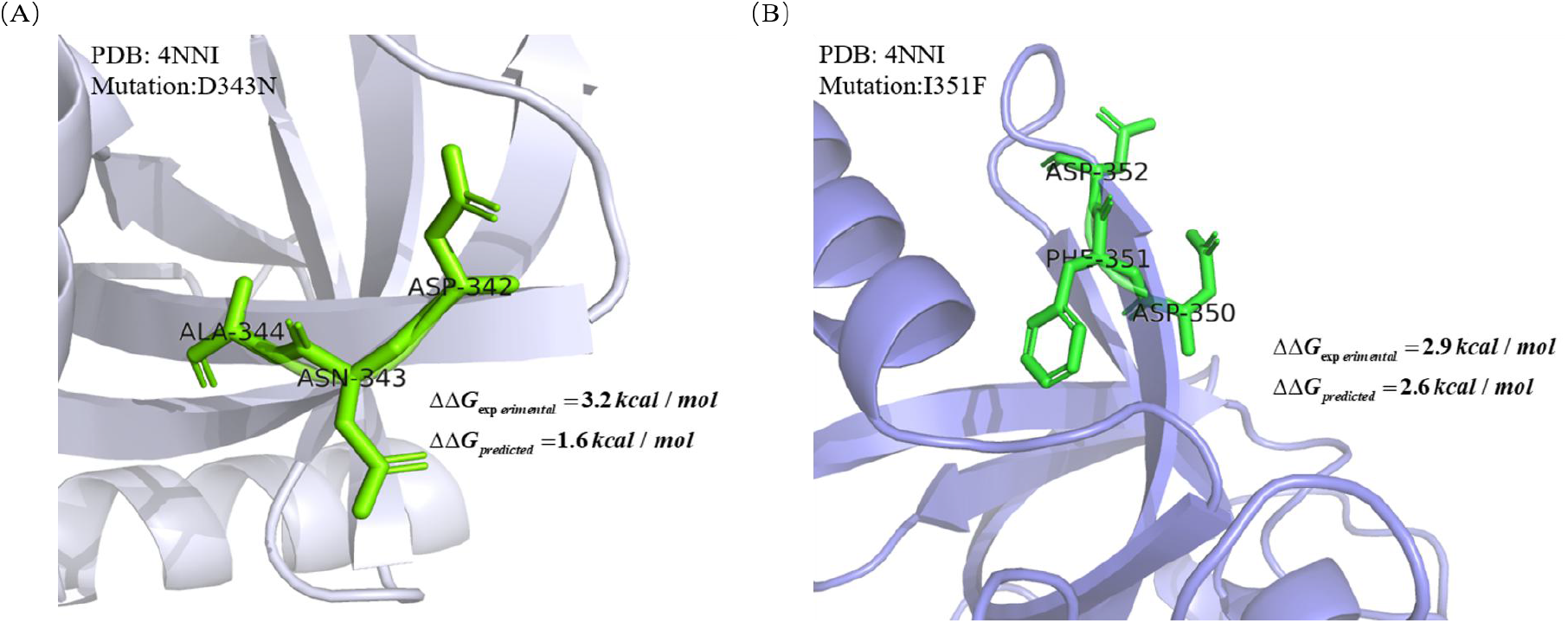
RpsA protein’s Gibbs free energy change prediction upon mutation and their residue local environment. (A) D343N mutation (B) I351F mutation

### 4.4 Time consumption study

Since we only extracted the information from the mutant residue and its two neighbors, the strategy substantially reduces the training and testing time by one or two orders of magnitude.

For example, the training time for the S2648 data set is down to 6-10 seconds per epoch after mutation part extraction(Table 2). Even if trained for 400 epoches, it only takes around one hour. Plus, the testing last for only 1 second. This highly efficient manner clearly caters the high throughput requirement in the pharmaceutical industry, where high volume data needs to be tested. It is noteworthy that removing the redundant data also boosts the overall prediction accuracy as the irrelevant information has been eliminated.

**Table 2.** Time consumption comparison between with and without mutant part data extraction.

|  | data extraction | w/o data extraction |
| --- | --- | --- |
| training time per epoch | 6~10 seconds | 900~1000 seconds |
| testing time | 1 seconds | 15~20 seconds |

## 5 Conclusion

Here we pioneered into predicting protein stability change upon mutation with an end-to-end GNN. To exclude irrelevant features and speed up the training, we first extracted the mutant part data. Subsequently, the model leveraged a gated GNN to capture the molecular features by message passing. In addition, it incorporated the raw molecular coordinates into the framework to predict ΔΔG. Rigorous experimental evaluations show that our model performed highly competitively on the S2648, S3214, S^sym^, and Myoglobin datasets. Furthermore, the method led to reasonable estimation for clinical drug resistance prediction. The substantial success over the task suggests a new strategy for swift protein stability change prediction and enlightens future GNN based method for improvement.

